# The Narrowing of Dendrite Branches across Nodes follows a well-defined Scaling Law

**DOI:** 10.1101/2020.04.13.039388

**Authors:** Maijia Liao, Jonathon Howard

## Abstract

The systematic variation of diameters in branched networks has tantalized biologists since the discovery of da Vinci’s rule for trees. Da Vinci’s rule can be formulated as a power law with exponent two: the square of the mother branch’s diameter is equal to the sum of the squares of those of the daughters. Power laws, with different exponents, have been proposed for branching in circulatory systems and in neurons. The laws have been derived theoretically, based on optimality arguments, but, for the most part, have not been tested rigorously. In the case of neuronal dendrites, diameter changes across branch points have functional implications for the spread of electrical signals: for example, Rall’s law with an exponent of 3/2 maximizes propagation speeds of action potentials across branch points. Using a super-resolution method to measure the diameters of all dendrites in highly branched *Drosophila* Class IV sensory neurons, we have tested Rall’s law and shown it to be false. In its place, we have discovered a new diameter-scaling law: the cross-sectional area is proportional to the number of dendrite tips supported by the branch plus a constant, corresponding to a minimum dendrite diameter. The law accords with microtubules providing force and transport for dendrite tip growth. That the observed scaling differs from Rall’s law suggests that constraints imposed by cell biological mechanisms may impact electrical signaling in neurons. Our new scaling law generalizes to other branched processes such as the vasculature of plants and the circulatory system of animals.

**Significance Statement:** To study the systematic variation of dendrite diameters, we have established a super-resolution method that allows us to resolve dendrite diameters in *Drosophila* Class IV dendritic arborization neurons, a model cell for studying branching morphogenesis. Interestingly, they do not follow any of the known scaling laws. We propose a new scaling law that follows from two concepts: there is an incremental cross-sectional area needed to support each terminal branch, and there is a minimum branch diameter. The law is consistent with dendrite growing by tip extension and being supported by microtubule-based transport. If the law generalizes to other neurons, it may facilitate segmentation in connectomic studies.

## Introduction

Branched networks are ubiquitous in nature, ranging in size from watercourses and trees to cellular organelles and the cytoskeleton (1–6). Often, branch diameters change systematically throughout the network, with proximal branches thicker than distal ones. This variation is usually interpreted as an adaptation to, or consequence of, the flow of materials and/or information through the network (7, 8). To describe the changes in diameter over branch points, allometric (or scaling) relations of the form

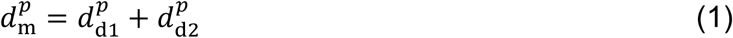

have been proposed, based mainly on theoretical arguments (see below), where *d*_m_ (*d*_d1_, *d*_d2_) is the mother (daughters) diameter and *p* is the exponent. Among the most well-known laws are da Vinci’s rule for trees (*p* = 2) (9), Murray’s law for vascular and pulmonary systems (*p* = 3) (7), and Rall’s law for neuronal processes (*p* = 3/2) (10). In this work, we ask whether neuronal dendrites obey these or other scaling laws?

Scaling laws have been derived theoretically using optimality arguments. For example, Murray’s law for the vasculature minimizes the frictional dissipation associated with moving fluid through pipes, given that the volume of the blood is constant (7). Rall’s law minimizes the propagation time of action potentials (11) and the decrement of graded electrical signals across dendrite branch points (12) (see Supplemental Information, Figure S1); it assumes that the density of ion channels is constant and the cost of building or maintaining the surface area of the dendrite is minimized. Da Vinci’s law has been reformulated as the “pipe model” for plants, in which a fixed cross-section of stems and branches is required to support each unit amount of leaves (13). Experimental support for scaling laws, however, is scarce because of the difficulties of imaging entire branched networks and because intrinsic anatomical variability may obscure precise laws (14). Thus, it is an open question whether scaling laws such as Equation 1 apply in biological systems.

To quantitatively test diameter-scaling laws in neurons, it is necessary to study cells in which all the dendrite diameters can be measured. These criteria are satisfied by *Drosophila* larval Class IV dendrite arborization neurons, which serve as a model system for studying dendrite morphogenesis (15). These neurons, which have up to 2000 branches, form an approximately planar network array (16) that tiles the external surface of the larvae like chain mail (17) (Figure 1a-b) and function as sensory receptors of nociceptive (18) and proprioceptive (19) inputs. They can be marked with a GFP-tagged membrane protein expressed under a cell-specific promoter (20). We developed a super-resolution method to measure even the finest dendrite diameters in these cells and have discovered a novel diameter-scaling law that holds throughout development.

**Figure 1.**
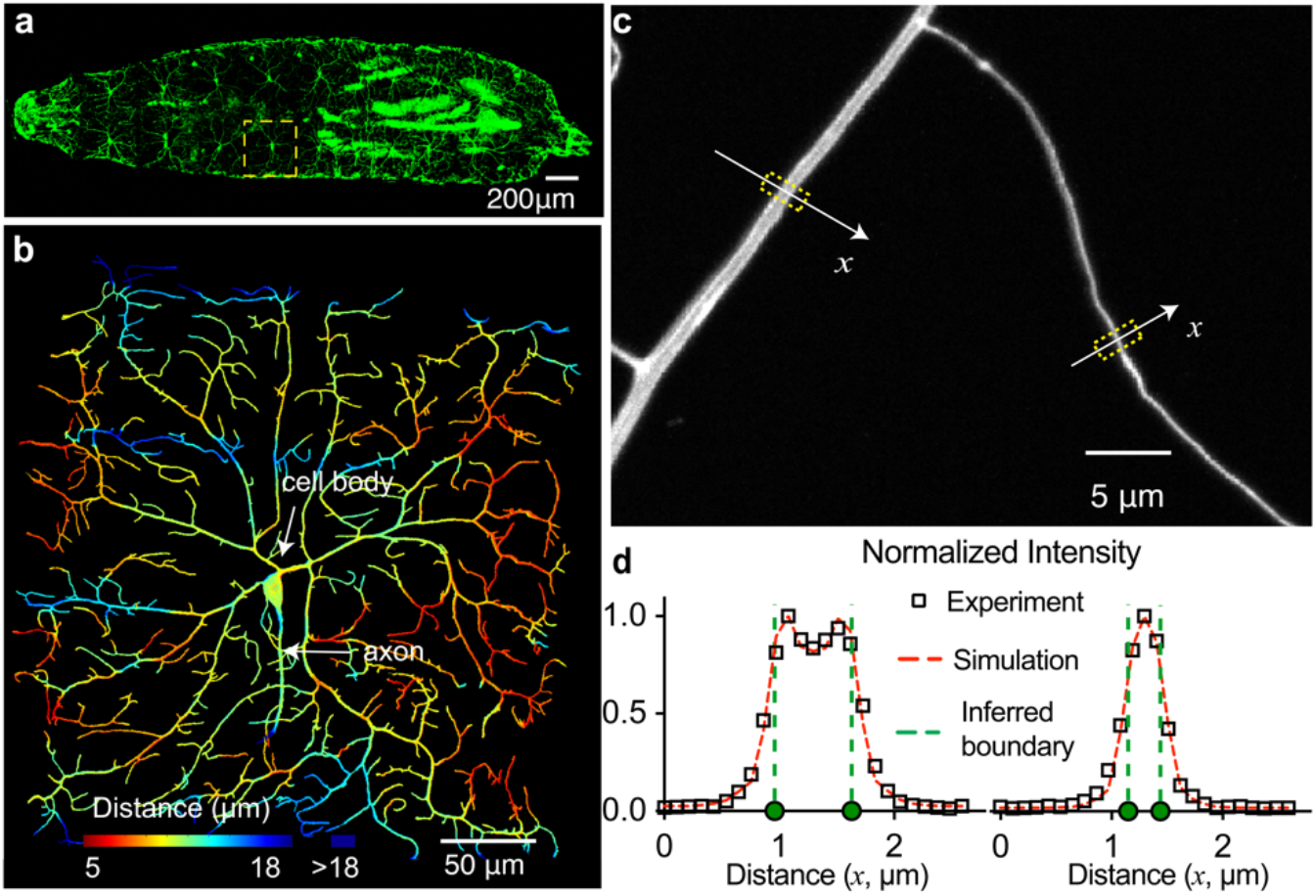
Precise measurement of dendrite branch diameter. **a** Third instar larva expressing GFP in its Class IV neurons. One cell in abdominal segment 3 (A3) is boxed. Anterior is left. **b** Maximum projection image of a Class IV neuron from the A3 segment constructed from 76 stacks (separated by 0.2 μm) from a spinning disk confocal microscope (Nikon spinning disk, 50 μm pinholes, 40X water-immersion objective with numerical aperture (NA) equal to 1.25) of a 92 hr after-egg-lay larva (*Shi*^*ts1*^*;;ppkCD4-tdGFP*). The distance of dendrites from the coverslip is color-coded. Note that the cells are mostly within a depth of ~12 μm, about 4% of their width. This dendritic tree has three arbors emerging from its cell body. **c** Maximum-intensity-projection images of a thick and a thin dendrite of a ~100-hr larva expressing tdGFP using a 60X WI objective with NA = 1.2. The yellow boxes with width 1.1 μm indicate the regions of the dendrite where the intensity profiles in **d** were obtained. **d** Intensity profiles of the middle section of the confocal stack for the thick and thin dendrites in **c**. The inferred diameters from eight 1000-step simulations are 667 ± 9 nm and 292 ± 11 nm (mean ± SD). Mean values are indicated by green dashed lines.

## Results

### Dendrite branch diameters are precisely measured using a super-resolution method

To test scaling laws, accurate measurements of dendrite branch diameters are essential. Conventional image analysis techniques based on thresholding techniques (14, 21, 22), cannot resolve the diameters of the finest dendrites (<300 nm). To accurately measure dendrite diameters, we developed a super-resolution method (Figure S2, Materials and Methods) that uses Monte-Carlo optimization to fit model images to experimental images obtained by spinning-disk confocal microscopy of Class IV dendrites whose surfaces were labeled with a membrane-associated fluorescent protein (Figure 1c). The model images are hollow cylinders with randomly distributed fluorophores on their surfaces convolved with the point spread function of the confocal microscope. The experimental images are *z*-stacks. The principle behind the method is that even for the thinnest dendrites, a best fit to the experimental data is obtained using a cylinder of non-zero diameter (i.e. the cylinder gives a better fit to the data than a line). Even though the lateral resolution for the confocal microscope is ~200 nm (≈ 0.5*λ*/NA, where NA = 1.2 is the numerical aperture of the water immersion objective and *λ* ≈ 520 nm is the emission wavelength), the method can resolve diameters of model dendrites below 150 nm (Table 1), which is less than the diameter of the finest dendrites in these cells. Figure 1d shows that our simulated intensity profiles are close to the ones measured in Class IV cells. An additional validation of our method based on dendrite intensities is described below. Thus, the diameters of even the thinnest dendrites can be measured.

**Table 1.** Accuracies of diameter measurements.

|  |  |  |  |  |  |
| --- | --- | --- | --- | --- | --- |
| True diameters of the model<br>image (nm) | 144 | 216 | 432 | 648 | 864 |
| Measured diameters<br>(Mean ± SD nm, $N = 18$<br>simulations) | 140±25 | 222±16 | 422±24 | 659±22 | 880±19 |

The fluorophore density 1646/ $\mu\text{m}^2$ was chosen based on estimations from the absolute intensity. Average over three cylinder orientations: 0.0, 0.5 and 1.0 rad relative to axial direction.

### Dendrite branch diameters do not obey known scaling laws

Before testing scaling laws, we first confirmed that dendrite branches have well-defined diameters. We imaged one to three arbors of dendrite trees from six fully developed third instar larvae (~132 hours after the time the egg was laid; egg lay was defined as time 0). We measured diameters, defined in Figure 2a, at three positions along 306 branches from the six larvae, avoiding locations where the membrane bulged due to the presence of membrane-bounded organelles. We plotted the normalized diameter against the normalized displacement along the branch, where zero displacement is towards the cell body (Figure 2b, Materials and Methods). The slope was −2.3%, which is much smaller than the average diameter from the proximal, mother branch (−18%) and to the distal, daughter branches (−11%) (Figure 2b). Thus, branch segments have approximately constant diameters, and diameters change primarily across branch nodes.

**Figure 2.**
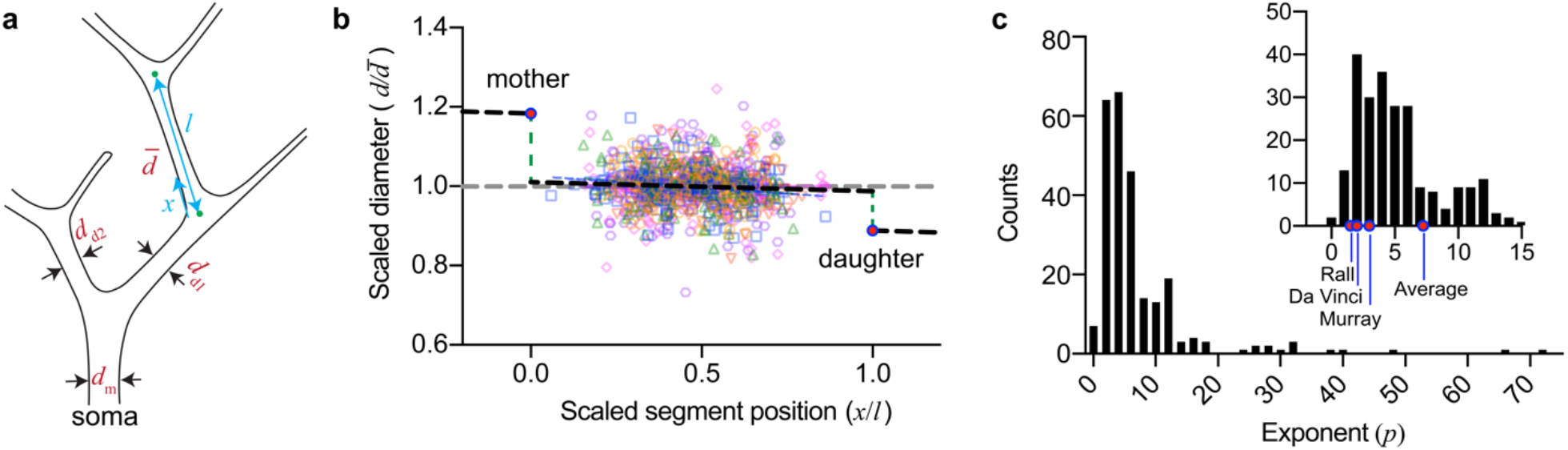
Dendrite branch diameters do not obey existing scaling laws. **a** Definitions of diameters and branch length. **b** Scaled diameters (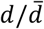 where 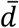 is the mean diameter of the branch) plotted against scaled position along the branch (*x*/*l* where *l* is the total length of the branch). Two branches were selected for each branch order between 0-34 for 6 neurons from six 132-hr larvae. The number of diameter measurements from each neuron ranged from 90 to 204. The black dashed line represents the linear regression with slope −0.023 ± 0.013 (± SE, *P* = 0.078 using a two-sided *t*-test). The left and right magenta filled circles are the mean diameters of the mothers and the daughters. **c** Histogram of values of the exponent *p* for 253 dendrite branch points from the six 132-hr larvae. The average exponent is 7.2 and the standard deviation is 8.8. Inset shows an enlarged view with magenta filled circles indicating exponents predicted by the Rall, da Vinci and Murray scaling laws.

To test the scaling law of Eq. (1), we calculated the exponents, *p*, for 253 branch points in six cells from the six 132-hr larvae that were used in Figure 2b. These branch points sampled the full range of branch orders. The mean exponent of 7.2 (Figure 2c) does not support any of the aforementioned scaling laws (i.e. Rall, da Vinci, Murray). Furthermore, the wide range of exponents (SD = 8.8) is not consistent with any specific scaling law. Alternative allometric relations are therefore required.

### The systematic change of dendrite diameters can be described by a new scaling law

Among a wide range of allometric relations that we tested, we discovered an approximately linear relationship between the leaf number, *n*, of a branch (the number of tips that the branch supports, Figure 3a, Figure S3) and the branch’s cross-sectional area (*A*) : *A* = *βn* + *A*_0_, where *β* is the slope and *A*_0_ is the *y*-intercept (Figure 3b). Because the leaf number in a bifurcating network satisfies the simple allometric relation: *n*_m_ = *n*_d1_ + *n*_d2_ where *n*_m_ (*n*_d1_, *n*_d2_) are the leaf numbers of the mother (daughters), the linear relationship implies a new allometric scaling law:

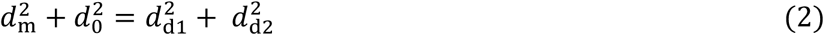

where 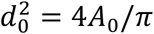 (see Supplemental Information for derivation). The interpretation of *d*_0_ is that there is a minimum dendrite diameter: the terminal branch with one tip has a diameter 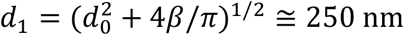.

**Figure 3.**
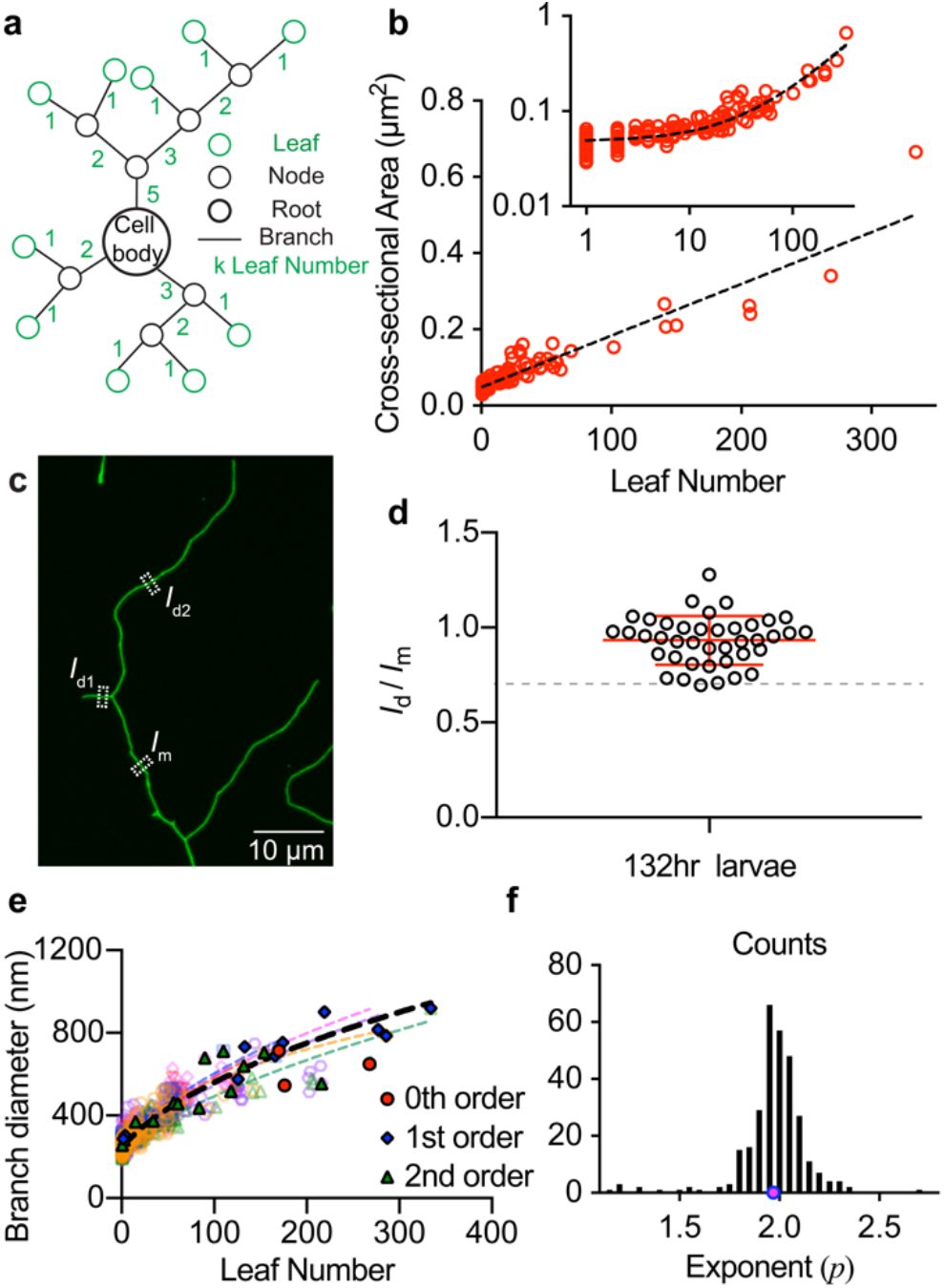
A new scaling law for bifurcating dendrites. **a** Definition of the leaf number of a bifurcating tree. **b** Cross-sectional area vs. leaf number for one 132-hr larva. Black dashed line indicates linear regression with slope *β* 1.36 ± 0.03 × 10^3^ nm^2^ and *y*-intercept 4.8 ± 0.1 × 10 nm^2^. Inset: log-log plot. **c** Maximum projection image of a bifurcation which has two terminal, daughter branches labeled. The white boxed regions with lateral width 1.1 μm indicate the regions of dendrite where the overall intensity *I*_m_(*I*_d1_, *I*_d2_) for mother and daughters branches of 132-hr larva were obtained]. **d** Intensity ratios of terminal daughters to the mother for 20 bifurcations from four 132-hr larvae. Red lines 0.93 ± 0.13 indicate mean and standard deviation. **e** Branch diameter vs. leaf number for six 132-hr larvae. Colored dashed curves indicate the non-linear fits to 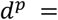 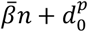 for each neuron. Black dashed curve shows fitting for all data points: *p* = 2.05 ± 0.06, *d*_0_ = 244 ± 3 nm (± SE, 1389 data points). Filled symbols indicate 0^th^, 1^st^ and 2^nd^ branch orders, where the 0^th^ order branch originates in the cell body. **f** The histogram shows the distribution of values of the exponent *p* fitted from: 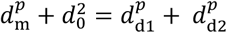 for 303 neuron branch points from six 132-hr larvae. The average exponent value, denoted by magenta circle, is 1.97 ± 0.17 (± SD).

A minimum dendrite diameter is not due to our failure to resolve the diameters of the finest dendrites. First, we note that the *y*-intercept is Figure 3b is significantly greater than zero (*p* ≪ 0.005, student’s t-test using the mean and SE in the legend to Figure 3), showing that our data strongly support the existence of a minimum diameter. Furthermore, the intensities of the terminal branches are similar to those of their mothers (Figure 3c,d), consistent with terminal branches (leaf number 1) having the same diameters as their mothers (leaf number 2). If there were no minimum diameter and the scaling law with exponent 2 continued across the most distal branch point, we would expect the daughter intensity to be 1/√2 ≅ 70% that of the mother (dashed line in Figure 3d). However, the measured intensities of the terminal branches are close to those of their mothers: 93% ± 13% (mean ± SD, *N* = 20, Figure 3d). Note that if there were less transport of fluorescent protein into the distal branch, we would expect a further reduction in intensity below 70%, contrary to what is observed. Thus, the minimum diameter in the scaling law is well supported experimentally and not likely to be due to a failure of the super-resolution method. The small scatter in the diameter measurements, with a coefficient of variation of 0.14 ≅ 0.13/0.93, confirms that the precision of the diameter measurement is high.

To test whether the exponent of 2, which corresponds to cross-sectional area, is the most appropriate, we fit the experimental data from the six 132-hour larvae to 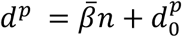. This scaling law provided a good fit (Figure 3e) and the exponent *p* was narrowly distributed with an average of 1.97, close to that for cross-sectional area (Figure 3f). Thus, our new scaling law with a minimum diameter and an area dependence provides a good description to diameter scaling in Class IV dendrites.

### Scaling is invariant over development

Class IV neurons grow throughout larval development from a cell width of ~60 μm at 24 hrs to ~400 μm at 132 hrs (Figure S4). To complete the description of allometric scaling, we applied our scaling relation 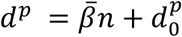 over all stages of development. Figures 4a-c show that the scaling parameters are roughly constant throughout development. The average value over developmental time of the exponent *p* is 2.40 ± 0.80, (mean ± SD, 31 larvae), corresponding approximately to area scaling (Figure 4a). The average value over developmental time of the cross-sectional area per leaf is 1970 ± 670 nm^2^ (± SD, 31 larvae) (Figure 4b). The average value of the terminal diameter 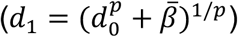 is 237 ± 14 nm (±SD, 31 larvae) (Figure 4c), and is independent of developmental time (Figure S5). Thus, the scaling law holds throughout the development of Class IV cells. While there is substantial variability in the exponent for over larva development, the exponent is more tightly centered around *p* = 2 in the older larvae (Figure 4a) ; this tightening of the distribution is evidence for a maturation process that adjusts diameters over development.

**Figure 4.**
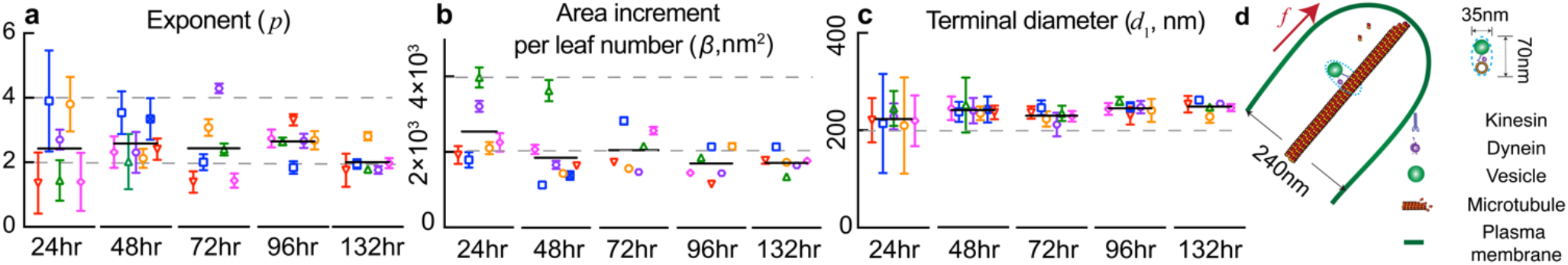
New scaling law measured at different developmental stages. **a** Scaling exponent *p*; **b** increment per leaf, *β*; **c** terminal diameter, *d*_1_; measured at different developmental stages (hrs after egg lay). Symbols indicate means and vertical lines indicate SEs obtained from fits to six or seven individual cells at each stage of 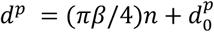. Black lines denote averages. **d** Sizes of the dendrite tips relative to microtubules and motors. Tip of a Class IV neuron with a microtubule. Each microtubule and vesicle occupy a cross-sectional area of 35 × 75 nm^2^, corresponding to the incremental area per leaf. The microtubule has diameter 25 nm.

## Discussion

Our data show that scaling laws of the form Eq. (1), which includes Rall’s law, do not hold for *Drosophila* Class IV neurons: there is no fixed exponent that accounts for all the bifurcations. The reason for the variability in exponents is that the branches in Class IV cells have a minimum diameter, according to Eq. (2), so that the daughters in more distal branches have similar diameters to their mothers and therefore a higher apparent exponent. For more proximal dendrites, by contrast, we found that the exponent approaches 2. As a consequence, there is a broad range of exponents, the great majority of which exceed the Rall exponent of 3/2. This finding agrees with studies in crustacean somatogastric ganglion neurons (14) and mammalian dendrites and axons (11, 23) in which the exponents varied from 0.5 to 4 (95% range), with medians between 2 and 3.

The deviation from Rall’s law has potential functional consequences. If the conduction speed of the action potentials (24) in these unmyelinated dendrites (16) is maximized across branch points, then there must be a compensatory change in ion channel density: the daughter density must be lower than that of the mother in the thinnest branches and higher in the thickest branches. Thus, the failure of scaling laws, and in particular Rall’s law, suggests that there may be a trade-off between cell geometry and molecular localization.

Class IV data fits a new scaling law (Eq. (2) with *p* ≅ 2): the area increases in proportion to the increase in the number of dendrite tips and there is a minimum diameter of terminal branches of 240 nm. We hypothesize that the new scaling law arises from cell biological and developmental constraints. The increase in cross-sectional area (*p* = 2) with leaf number can be interpreted as nutrient flux through the cytoplasm being proportional to the rate of elongation of dendrites, which is proportional to the number of dendrite tips. Class IV neurons grow throughout development and the primary mechanism by which dendrite length increases is through tip extension (ref. (25) and manuscript in preparation). This interpretation assumes that dendrite transport velocity is independent of dendrite diameter, as expected if motor-driven transport has a constant speed, so that flux is proportional to cross-sectional area. The extra cross-sectional area per tip of 2000 nm^2^ is similar to the area occupied by one microtubule and a 30-nm vesicle carried along it by a motor protein (an ellipsoid with axes 35 nm and 70 nm) (Figure 4). A similar argument has been used to estimate the minimum diameter of axons (26). An additional microtubule may therefore be required in a supporting branch to convey the materials and signaling molecules towards and away from each additional tip. Thus, diameter scaling may be a consequence of the metabolic needs of the cells, supporting the view that energetics is an important design consideration in the nervous system (27).

The observed minimum diameter of 240 nm in Class IV cells may be a consequence of the force required for dendrite extension. To extend a tube of membrane from a low-curvature bilayer, a force, *f*, is required to bend the membrane into a cylinder: *f* = 4*πκ*/*d* where *κ* is bending stiffness and *d* is the cylinder’s diameter (28). Therefore, the smaller the diameter, the larger the force. For a membrane stiffness of 40 pN/nm (29), a diameter of 240 nm requires a force of ≅ 2 pN (Figure 4d), which is similar to the force generated by a single growing microtubule (30) or a single microtubule-based motor (31) driving microtubule sliding (32). The force will be even higher if other structures, such as internal membranes or cytoskeleton, are also strained during dendrite extension. Thus, a dendrite of diameter 240 nm could be extended by a single motor or microtubule, but a narrower dendrite would require the concerted action of multiple microtubules and/or motors.

Though these arguments are conjectural, they demonstrate the plausibility that dendrite geometry is limited by cell biological constraints. Specifically, these arguments are consistent with microtubules being critical for providing force and materials necessary for dendrite growth (33, 34). Taken together, our data and analysis support Cajal’s conjecture that “all of the morphological features displayed by neurons appear to obey precise rules [e.g. scaling laws] that are accompanied by useful consequences [e.g. optimizing transport required for growth]” (35). Should our new scaling law generalize to other dendrites, it is expected to facilitate segmentation in connectomic studies (36–38) because knowing the rules for dendrite morphology will provide priors that constrain connectomic maps, just as Ramachandran plots constrain and validate protein structures.

In addition to neuronal systems, our new scaling law may generalize to other biological systems. The pipe model for plants, which is equivalent to da Vinci’s rule, posits that the amount of leaves and fruit is proportional to the cross-sectional area of soft (living) wood (13, 39). Including a minimum diameter provides a better fit to some of the experimental data (Figure 5a). We also applied the new scaling law to a reconstructed pig arteriolar tree (21), and, again, including a minimum area provides a better fit (Figure 5b). The minimum diameter of the distal branches estimated from Figure 5b is 6.2 μm, close to 6-μm diameter of pig red blood cells (40). Thus, scaling laws that consider tubular networks having minimum diameters may apply generally to branching in a broad range of biological networks.

**Figure 5.**
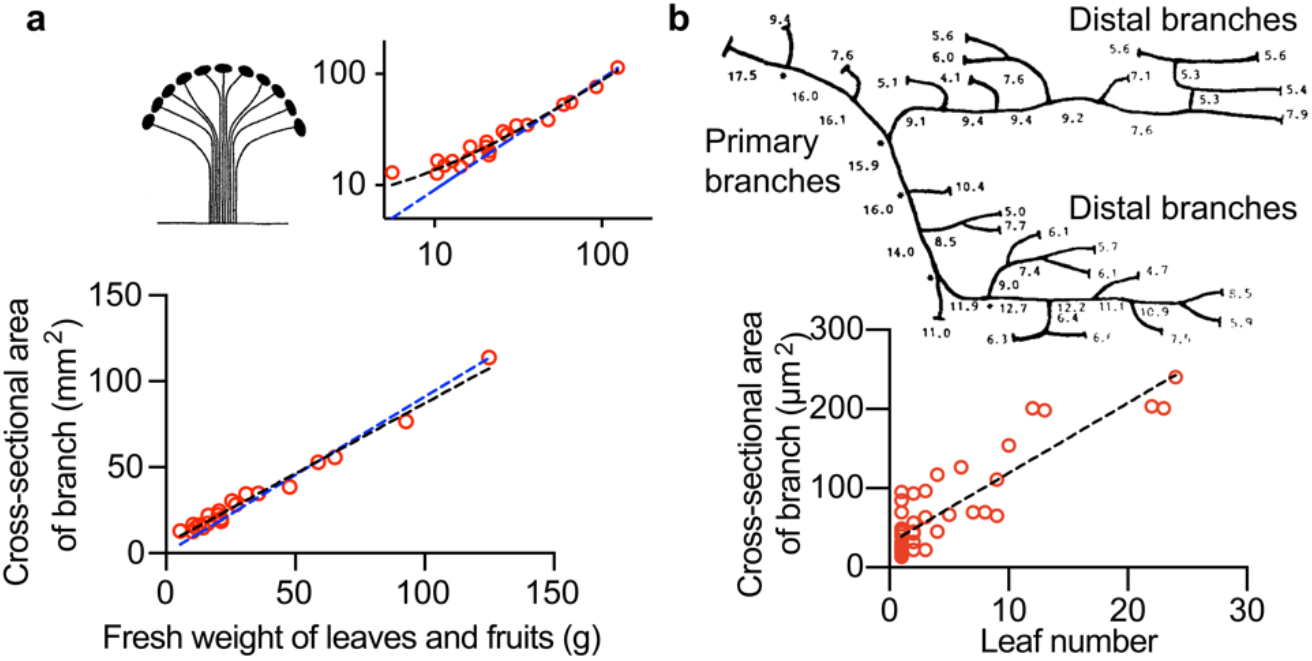
Scaling of tree branches and vasculature accords with the new scaling law. **a** Linear relation between the area of the cross-section of a branch of *Ficus erecta* and the weight of leaves and fruits born by that branch. Black dashed line indicates linear regression with slope 0.82 ± 0.02 mm^2^ · g^−1^ and *y*-intercept 5.51 ± 1.08 mm^2^. Blue dashed line indicates linear regression fitting with slope 0.91 ± 0.02 mm^2^ · g^−1^ and *y*-intercept is set as 0. Data points were digitalized from ref. (13). A non-zero y-intercept provides a better fit. The insets show log-log scale plot and the idea behind the pipe model, which is equivalent to da Vinci’s rule. **b** Upper panel. Schematic diagram of a reconstructed arteriolar tree adapted from Fig. 2 of ref.(21). Numbers indicate diameters of each vessel segment. Primary and distal branches can be distinguished from branch diameters. Lower panel. Branch cross-sectional areas are plotted versus leaf number. Black dashed line indicates linear regression fitting with slope *β* 8.86 ± 0.69 *μ*m^2^ and *y*-intercept 30.5 ± 5.0 μm^2^. Including a minimum diameter provides a better fit to the mother-daughter ratio distribution (Fig. S7).

## Materials and Methods

### *Drosophila melanogaster* strains

The Shibire strain *Shi*^*ts1*^;; ppk-CD4-tdGFP was generously provided by Fernando Vonhoff. *Shi*^*ts1*^ is a temperature sensitive mutation that immobilizes the larva when the temperature is raised from 25.0 °*C* (the permissive temperature) to 30.5 °*C* (the non-permissive temperature). These mutants have normal development at the permissive temperature.

### Spinning Disk Confocal Imaging

Embryos were collected for 2 hrs on apple juice agar plates with a dollop of yeast paste and aged at 25 °C in a moist chamber. The plates containing the first batch of embryos were discarded as the dendrite morphology of Class IV neurons is less consistent in those animals (16). Larvae were immobilized individually on agarose pads (thickness 0.3-0.5mm) sandwiched between a slide and a coverslip. The imaging was done using a spinning disk microscope: the Yokogawa CSU-W1 disk (pinhole size 50 μm) built on a fully automated Nikon TI inverted microscope with perfect focus system, an sCMOS camera (Zyla 4.2 plus sCMOS), and running Nikon Elements software. The Shibire larvae were paralyzed at 30.5 °C during imaging. Individual neuron image stacks used for super-resolution measurements were acquired with a 60X 1.2 NA oil lens with a *z* step size 0.16μm. The whole-larva images were acquired with a 4X 0.2 NA objective. To ensure the neuronal morphology was not altered at the non-permissive temperature, each image acquisition process was completed within 20 min. Class IV neurons in A3–A5 segments from six to seven larvae were collected at each developmental stage.

### Model Image Construction

Our generative model aims to be an accurate physical description of the microscope imaging. Creating this model requires a detailed understanding of image formation in the confocal microscope. The sample is illuminated with a laser focused through the objective lens. The objective captures the emitted light from fluorophores and focuses it through a pinhole to reject out-of-focus light. Based on the physical setup, we can describe the confocal image using four main generative model components (41, 42). (i) Platonic image *Π*(*x*) – the fluorophores randomly distributed on a cylinder surface, with pixel size 9 nm. The fluorophore density, on the order of 1000 μm^−2^, is based on estimations of the single-fluorophore intensity in the spinning-disk microscope. (ii) Illumination field *I*(*x*) – the light intensity as a function of position. The illumination field is described as a product of an *x*-*y* illumination and a *z* modulation: *I*(*x*) = *I*_xy_(*x*, *y*) × *I*_z_(*z*). In practice, aberrations due to refractive index mismatches cause a dimming of the illumination with depth into the sample (43). Since this overall dimming only depends on the depth *z* from the interface and not on the *x*-*y* position in the sample, it is natural to describe the illumination field as a product of a uniform *x*-*y* illumination and a *z* modulation. (iii) Point spread function *P*_conf_(***x***, ***x***′) – the image of an individual fluorophore due to diffraction of light. The scalar Gibson and Lanni model (44, 45) was used; it gave similar results to the vectoral-based PSF model (46). The pinhole size of the confocal microscope was also incorporated into *P*_conf_(***x***, ***x***′) (47, 48). (iv) Image noise *B*(***x***) – autofluorescent signal from the surrounding environment. These components are combined to form the image through convolution: *M*(***x***) = *B*(***x***) + ∫ *d*^3^*x*′*I*(***x***′)*II*(***x***′)*P*_conf_(***x*** − ***x***′; ***x***), which is sampled at discrete pixel locations to get the final image *M*(***x***) (in units of photon number). These photons are then transformed into a number of electrons based on quantum efficiency (QE), shot noise and the sCMOS parameters. Then the image is reduced to the desired camera resolution, for example, 108 nm/pixel. These values are fed to an electron-to-DN converter (digital number, considering the readout noise) to obtain model images (42) (Figure S2).

### Super-resolution method based on Monte-Carlo (MC) optimization

The super-resolution method generates simulated images *M*(***x***) described above. The tunable parameters of the optimization are: the amplitude of the spatially uniform background, hollow cylinder radius, sample refractive index, pinhole diameter, emission wavelength and cylinder orientation and relative position. At each step, a simulated image is generated based on the tuned parameters. Then a comparison between the simulated image and the model image is made. The MC optimization aims to find the radius that minimizes the square of the difference (49) : 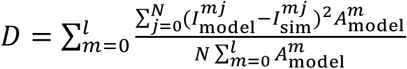, where *I*^*mj*^ stands for intensity profile across the cylinder as a function of pixel number *j* and image section *m* of the stack, *N* stands for the total number of pixels for image section *m*, and *A*^*m*^ stands for total pixel intensity of image section *m*. Subscript model stands for model images while sim stands for simulated images. Six 1000-step simulations were carried out for each branch diameter detection. The detected diameter is an average of results from six simulations.

### Analysis of Larvae Imaging Data

#### Class IV dendrite tracing

To quantify dendrite morphology, *z*-series images for a given neuron were projected onto a 2D image file and dendrite arbors were traced by tree toolbox software (50). To improve accuracy, the traces were further adjusted based on intensity profiles from maximum-intensity-projection images. Leaf number, branch order and Strahler number (see Figure S3) were measured from the traces.

#### Criteria for selecting dendrite branches for super-resolution analysis

Our super-resolution method aims to find diameters of hollow cylinders that best fit dendrite branches. We analyzed branches longer than 3 μm. We avoided regions containing endoplasmic reticulum (ER) or Golgi apparatus (51). These regions have higher intensities or larger diameters. Outliers of maximum intensity values and full widths at half maximum (FWHM) along the dendrite branches are first identified. The corresponding parts of branches are then removed. The remaining parts of the branches are then used for further analysis.

#### Scaled diameters and positions

After removing the parts of a branch that contain ER or Golgi apparatus, the remaining parts of that branch can be discontinuous. Through combining all remaining parts of the branch as a whole, we obtain a ‘new’ branch with all the above criteria satisfied. Then three positions are chosen at 1/3, 1/2 and 2/3 of the newly constructed branch and branch diameters are calculated at corresponding positions. The distance between the selected positions and the branch point nearest the soma in the original branch geometry is denoted as *x*_*i*_ (*i* = 1,2,3). Mean diameter of each branch is denoted by 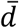 is an average of *d*_*i*_ (*i* = 1,2,3).

#### Exponent *p* measurements

Previous scaling laws have the form: 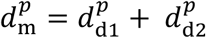. The exponent *p* is estimated by minimizing the difference between the right- and left-hand sides of the equation. To determine this value, we used the function *fsolve* in MATLAB.

## Data availability

Detailed materials and methods are reported in *SI Appendix*. The code to perform MC simulation will be available on GitHub upon publication (https://github.com/Maijia-cpu/Super-resolution-Method).

## Acknowledgments

We thank Xin Liang for initiating this project and members of the Howard laboratory for comments on an earlier draft. We benefited from discussions with Yuhai Tu (IBM Watson Laboratory, NY) and Howard lab members during the execution of this work, and comments on the manuscript from Rob Phillips. This study was supported by NIH grant DP1 MH110065.

## References

1. P. Ball, Branches: Nature’s patterns: a tapestry in three parts (OUP Oxford, 2009).

2. A. Rinaldo, R. Rigon, J. R. Banavar, A. Maritan, I. Rodriguez-Iturbe, Evolution and selection of river networks: Statics, dynamics, and complexity. Proc National Acad Sci 111, 2417–2424 (2014).

3. G. B. West, J. H. Brown, Life’s universal scaling laws. Physics today 57, 36–43 (2004).

4. S. M. Rafelski, et al., Mitochondrial network size scaling in budding yeast. Science (New York, N.Y.) 338, 822–824 (2012).

5. D. A. Fletcher, R. D. Mullins, Cell mechanics and the cytoskeleton. Nature 463, 485–92 (2010).

6. J. R. Banavar, A. Maritan, A. Rinaldo, Size and form in efficient transportation networks. Nature 399, 130–132 (1999).

7. C. D. Murray, The Physiological Principle of Minimum Work: I. The Vascular System and the Cost of Blood Volume. Proc National Acad Sci 12, 207–214 (1926).

8. A. Bejan, J. P. Zane, Design in nature: How the constructal law governs evolution in biology, physics, technology, and social organizations (Anchor, 2013).

9. J. P. Richter, The notebooks of Leonardo da Vinci (Courier Corporation, 1970).

10. W. Rall, Branching dendritic trees and motoneuron membrane resistivity. Experimental neurology 1, 491–527 (1959).

11. D. B. Chklovskii, A. Stepanyants, Power-law for axon diameters at branch point. Bmc Neurosci 4, 18 (2003).

12. Q. Wen, D. B. Chklovskii, A Cost–Benefit Analysis of Neuronal Morphology. J Neurophysiol 99, 2320–2328 (2008).

13. K. Shinozaki, K. Yoda, K. Hozumi, T. Kira, A quantitative analysis of plant form- the pipe model theory: I. basic analyses. Jpn J Ecol 14, 97–105 (1964).

14. A. G. Otopalik, et al., Sloppy morphological tuning in identified neurons of the crustacean stomatogastric ganglion. Elife 6, e22352 (2017).

15. Y.-N. Jan, L. Y. Jan, Branching out: mechanisms of dendritic arborization. Nat Rev Neurosci 11, 316–328 (2010).

16. C. Han, et al., Integrins Regulate Repulsion-Mediated Dendritic Patterning of Drosophila Sensory Neurons by Restricting Dendrites in a 2D Space. Neuron 73, 64–78 (2012).

17. W. B. Grueber, L. Y. Jan, Y. N. Jan, Tiling of the Drosophila epidermis by multidendritic sensory neurons. Dev Camb Engl 129, 2867–78 (2002).

18. R. Y. Hwang, et al., Nociceptive Neurons Protect Drosophila Larvae from Parasitoid Wasps. Curr Biol 17, 2105–2116 (2007).

19. W. Song, M. Onishi, L. Y. Jan, Y. N. Jan, Peripheral multidendritic sensory neurons are necessary for rhythmic locomotion behavior in Drosophila larvae. Proc National Acad Sci 104, 5199–5204 (2007).

20. W. B. Grueber, et al., Projections of Drosophila multidendritic neurons in the central nervous system: links with peripheral dendrite morphology. Development 134, 55–64 (2007).

21. G. S. Kassab, Y.-C. B. Fung, The pattern of coronary arteriolar bifurcations and the uniform shear hypothesis. Ann Biomed Eng 23, 13–20 (1995).

22. M. Zamir, J. A. Medeiros, Arterial branching in man and monkey. J Gen Physiology 79, 353–360 (1982).

23. C. Cherniak, M. Changizi, D. W. Kang, Large-scale optimization of neuron arbors. Phys Rev E 59, 6001–6009 (1999).

24. S.-I. Terada, et al., Neuronal processing of noxious thermal stimuli mediated by dendritic Ca(2+) influx in Drosophila somatosensory neurons. eLife 5, 618 (2016).

25. F. B. Gao, J. E. Brenman, L. Y. Jan, Y. N. Jan, Genes regulating dendritic outgrowth, branching, and routing in Drosophila. Genes & Development 13, 2549–2561 (1999).

26. A. A. Faisal, J. A. White, S. B. Laughlin, Ion-Channel Noise Places Limits on the Miniaturization of the Brain’s Wiring. Curr Biol 15, 1143–1149 (2005).

27. P. Sterling, S. Laughlin, Principles of neural design (MIT Press, 2015).

28. I. Derényi, F. Jülicher, J. Prost, Formation and Interaction of Membrane Tubes. Phys Rev Lett 88, 238101 (2002).

29. M. M. Kamal, D. Mills, M. Grzybek, J. Howard, Measurement of the membrane curvature preference of phospholipids reveals only weak coupling between lipid shape and leaflet curvature. P Natl Acad Sci Usa 106, 22245–50 (2009).

30. M. Dogterom, B. Yurke, Measurement of the Force-Velocity Relation for Growing Microtubules. Science 278, 856–860 (1997).

31. J. Howard, Mechanics of motor proteins and the cytoskeleton (Oxford Univ Press, New York) (2001).

32. W. Lu, P. Fox, M. Lakonishok, M. W. Davidson, V. I. Gelfand, Initial Neurite Outgrowth in Drosophila Neurons Is Driven by Kinesin-Powered Microtubule Sliding. Current Biology 23, 1018–1023 (2013).

33. P. W. Baas, A. N. Rao, A. J. Matamoros, L. Leo, Stability properties of neuronal microtubules. Cytoskeleton 73, 442–460 (2016).

34. L. C. Kapitein, C. C. Hoogenraad, Building the Neuronal Microtubule Cytoskeleton. Neuron 87, 492–506 (2015).

35. S. R. Cajal, Histology of the nervous system of man and vertebrates. History of Neuroscience (Oxford Univ Press, New York) (1995).

36. S. Seung, Connectome: How the brain’s wiring makes us who we are (HMH, 2012).

37. Y. Wang, A. Gupta, M. Toledo-Rodriguez, C. Z. Wu, H. Markram, Anatomical, Physiological, Molecular and Circuit Properties of Nest Basket Cells in the Developing Somatosensory Cortex. Cereb Cortex 12, 395–410 (2002).

38. C. S. Xu, et al., A Connectome of the Adult Drosophila Central Brain. Biorxiv, 2020.01.21.911859 (2020).

39. R. Lehnebach, R. Beyer, V. Letort, P. Heuret, The pipe model theory half a century on: a review. Ann Bot-london 121, 773–795 (2018).

40. R. Flindt, Amazing numbers in biology (Springer Science & Business Media, 2006).

41. M. Bierbaum, B. D. Leahy, A. A. Alemi, I. Cohen, J. P. Sethna, Light microscopy at maximal precision. Physical Review X 7, 041007 (2017).

42. D. Sage, et al., Quantitative evaluation of software packages for single-molecule localization microscopy. Nat Methods 12, 717–724 (2015).

43. S. Hell, G. Reiner, C. Cremer, E. H. Stelzer, Aberrations in confocal fluorescence microscopy induced by mismatches in refractive index. Journal of microscopy 169, 391–405 (1993).

44. S. F. Gibson, F. Lanni, Experimental test of an analytical model of aberration in an oil-immersion objective lens used in three-dimensional light microscopy. JOSA A 9, 154–166 (1992).

45. J. Li, F. Xue, T. Blu, Fast and accurate three-dimensional point spread function computation for fluorescence microscopy. J Opt Soc Am 34, 1029 (2017).

46. B. Richards, E. Wolf, Electromagnetic diffraction in optical systems, II. Structure of the image field in an aplanatic system. Proceedings of the Royal Society of London. Series A. Mathematical and Physical Sciences 253, 358–379 (1959).

47. S. Kimura, T. Wilson, Effect of axial pinhole displacement in confocal microscopes. Applied optics 32, 2257–2261 (1993).

48. T. Wilson, C. Sheppard, Theory and practice of scanning optical microscopy (Academic Press London, 1984).

49. F. Flicker, J. van Wezel, Charge order in NbSe2. Phys Rev B 94, 235135 (2016).

50. H. Cuntz, F. Forstner, A. Borst, M. Häusser, One Rule to Grow Them All: A General Theory of Neuronal Branching and Its Practical Application. Plos Comput Biol 6, e1000877 (2010).

51. M. M. Corty, B. J. Matthews, W. B. Grueber, Molecules and mechanisms of dendrite development in Drosophila. Development 136, 1049–1061 (2009).

