## Supplementary Information for "The Narrowing of Dendrite Branches across Nodes follows a well-defined Scaling Law"

**This PDF file includes:**

Supplementary text
Figures S1 to S7
SI References

### Supplementary Information

#### Rall's law — an optimization principle

For a (passive) cable of diameter $d$ with specific membrane conductance $g_{\mathrm{m}}$ (~1 $\Omega^{-1}\mathrm{m}^{-2}$) and specific resistivity of the internal medium $R_{\mathrm{i}}$ (~1 $\Omega\mathrm{m}$), the characteristic length constant is $\lambda = \frac{1}{2} d^{1/2}(g_{\mathrm{m}}R_{\mathrm{i}})^{-1/2}$ and the input impedance is $G_{\infty} = \left(\frac{\pi}{2}\right)(R_{\mathrm{i}}/g_{\mathrm{m}})^{-1/2} d^{3/2} \propto d^{3/2} {g_{\mathrm{m}}}^{1/2}$ (1). Rall discovered that if the diameters of the mother and daughter branches satisfy $d_{\mathrm{m}}^{3/2} = d_{\mathrm{d1}}^{3/2} + d_{\mathrm{d1}}^{3/2}$ and the channel density is constant, then the impedance is matched across branch points (in the sense that the mother's input impedance equals the sum of the daughters'), and the potential decreases exponentially with distance from the soma. This implies that each dendrite emerging from the soma contributes an electrical conductance corresponding to an infinite cylinder. In this way, he estimated the relative conductance of the soma and the dendrites.

*Minimization of signal loss across branch points* (2). The signal decrement across a branch point is $\Delta V = V_{\mathrm{d1}}(L_{\mathrm{d1}}/\lambda_{\mathrm{d1}} + L_{\mathrm{m}}/\lambda_{\mathrm{m}}) + V_{\mathrm{d2}}(L_{\mathrm{d2}}/\lambda_{\mathrm{d2}} + L_{\mathrm{m}}/\lambda_{\mathrm{m}}) \propto (V_{\mathrm{d1}} + V_{\mathrm{d2}})L_{\mathrm{m}}/\sqrt{d_{\mathrm{m}}} + V_{\mathrm{d1}}L_{\mathrm{d1}}/\sqrt{d_{\mathrm{d1}}} + V_{\mathrm{d2}}L_{\mathrm{d2}}/\sqrt{d_{\mathrm{d2}}}$ where $V_{\mathrm{d1}}$ ($V_{\mathrm{d2}}$) is the signal in daughter 1 (daughter 2) and we assume that the lengths of the mother and daughter branches ($L_{\mathrm{m}}$, $L_{\mathrm{d1}}$ and $L_{\mathrm{d2}}$) are much shorter than the length constants. If the signal decrement is a minimum for a fixed total surface area then a relation between the mother and daughter diameters can be determined by minimizing $\Delta V + \alpha(d_{\mathrm{m}}L_{\mathrm{m}} + d_{\mathrm{d1}}L_{\mathrm{d1}} + d_{\mathrm{d2}}L_{\mathrm{d2}})$, with respect to $d_{\mathrm{m}}$, $d_{\mathrm{d1}}$ and $d_{\mathrm{d2}}$. $\alpha$ is a Lagrange multiplier and the bracketed term is proportional to the total surface area of the branch. Minimization gives $d_{\mathrm{m}}^{3/2} = d_{\mathrm{d1}}^{3/2} + d_{\mathrm{d2}}^{3/2}$, termed Rall's law to distinguish it from Rall's original relationship, which had no functional interpretation, being instead a computational tool for calculating dendritic resistances. The optimal ratio of daughter branches diameters depends on the relative signals the branches carry: $d_{\mathrm{d1}}/d_{\mathrm{d2}} = (V_{\mathrm{d1}}/V_{\mathrm{d2}})^{2/3}$. If the signals are the same in the two dendrites, then $d_{d1} = d_{d2}$. A fixed total surface area is equivalent to there being a cost of building or maintaining a dendrite that is dominated by the surface area.

*Minimization of signal delay loss across branch points*. A similar argument can be made for minimizing action potential delays across branch points. If the membrane impedance is constant (e.g. constant channel densities), then, for unmyelinated fibers, the propagation speed is proportional to the square root of dendrite diameter ($d^{1/2}$) and the delay is proportional to $d^{-1/2}$. The total delay across a branch point if there are $w_{\mathrm{d1}}$ ($w_{\mathrm{d2}}$) action potentials in daughter 1 (daughter 2) is $\Delta T \propto (w_{\mathrm{d1}} + w_{\mathrm{d2}})/\sqrt{d_{\mathrm{m}}} + w_{\mathrm{d1}}/\sqrt{d_{\mathrm{d1}}} + w_{\mathrm{d2}}/\sqrt{d_{\mathrm{d2}}}$. If the total delay is a minimum for a fixed total surface area then a relation between the mother and daughter diameters can be determined by minimizing $\Delta T + \alpha(d_{\mathrm{m}} + d_{\mathrm{d1}} + d_{\mathrm{d2}})$ with respect to $d_{\mathrm{m}}$, $d_{\mathrm{d1}}$ and $d_{\mathrm{d2}}$. $\alpha$ is the Lagrange multiplier and the sum of the diameters is proportional to the surface area. We again recover Rall's law: $d_{\mathrm{m}}^{3/2} = d_{\mathrm{d1}}^{3/2} + d_{\mathrm{d2}}^{3/2}$. If the number of action potentials in the two dendrites are the same, then $d_{d1} = d_{d2}$.

*Volume constraint.* If the total volume is fixed, then Rall's law becomes: $d_{\mathrm{m}}^{5/2} = d_{\mathrm{d1}}^{5/2} + d_{\mathrm{d2}}^{5/2}$. If there are costs to both surface area and volume, then the 3/2 law will dominate at small diameters and the 5/2 law at large diameters.

*Myelinated fibers*. For myelinated fibers, the velocity is proportional to the diameter (3). In this case, the maximum velocity gives an exponent of 2 for surface area and 3 for volume (4).

**Leaf number is a reliable descriptor of branch diameter**

To look for new branching rules, it is necessary to enumerate the branches and nodes.

*(i) Branch order or depth* (Fig. S3a) describes the hierarchy of a tree measured from the root. Because the cell body can form three and sometimes four primary dendrites, we define the root of the arbors as the first bifurcation of the primary

dendrites. Thus, an entire dendrite is made up of three (or four) binary trees. The largest branch order is the arbor's height; because the trees are imperfect, each arbor can have different heights.

(ii) The *Strahler number* (5) describes the hierarchy starting from the tips (Fig.S3b). All terminal branches are assigned order 1. The remaining orders are then constructed in an iterative way: When two branches of order $k$ meet, the order of the parent branch is increased to order $k+1$. When two branches of different order meet, the higher order prevails.

(iii) The *leaf number* is defined in Fig. S3c.

To investigate how the widths of the branches depend on their position in the tree, we plotted the cross-sectional area against the branch order, Strahler number and the leaf number. Small branch orders can have both thick and thin dendrites, while high branch orders have thin dendrites (Fig. S3a). The spread of the data points at low orders reflects the asymmetry of branching and suggests that branch order is not a good numbering scheme for characterizing diameter variation. The cross-sectional area shows an increasing trend as the Strahler number increases (Fig. S3b), reflecting an increase in cross-section away from the distal tips. While the distal tips (Strahler number = 1) have a rather uniform cross-sectional area, the cross-sectional areas spread out as the Strahler number increases. This variability suggests that the Strahler number is also not a good descriptor of branch diameter. We then plotted the cross-sectional area against the leaf number (Fig.S3c and Fig. 3b). Compared to other numbering schemes, the leaf number leads to smaller scatter in the cross-sectional areas and the spread of data points remains roughly unchanged as the leaf number increases. Thus, leaf number is a reliable descriptor of branch diameter.

**New allometric scaling law derivation based on leaf number property**

Fig. 3b in the main text shows that there is an approximately linear relation between cross-sectional area and leaf number: $\pi d^2/4 = \beta n + \pi d_0^2/4$. Based on this observation, we wrote the relation in a more general way: $d^p = \bar{\beta} n + d_0^p$.

Because the leaf number satisfies the simple allometric relation: $n_{\mathrm{m}} = n_{\mathrm{d1}} + n_{\mathrm{d2}}$ where $n_{\mathrm{m}}$ $(n_{\mathrm{d1}}, n_{\mathrm{d2}})$ are the leaf numbers of the mother (daughters), we have: $(d_{\mathrm{m}}^{p} - d_{0}^{p})/\bar{\beta} = (d_{\mathrm{d1}}^{p} - d_{0}^{p})/\bar{\beta} + (d_{\mathrm{d1}}^{p} - d_{0}^{p})/\bar{\beta}$. Rearranging, we obtain the new allometric scaling law: $d_{\mathrm{m}}^{p} + d_{0}^{p} = d_{\mathrm{d1}}^{p} + d_{\mathrm{d1}}^{p}$.

**Similarity exists between pipe model and our new scaling law**

The pipe model for plants, which is equivalent to da Vinci's rule, posits that the amount of leaves is proportional to the cross-sectional area of soft (living) wood. Shinozaki *et al.*(6) discovered a linear relation between cross-sectional area of branches of *Ficus erecta* and total amounts of leaves and fruits supported by those branches (Fig. 5a). Similarly, in Class IV dendritic branches, a linear relation between cross-sectional area and leaf number is also observed. The linear relation can be well described by $\pi d^2/4 = \beta n + \pi d_0^2/4$ (Fig. 3b) considering the minimal cross-sectional area. In Shinozaki *et al.* 's paper, however, the minimal cross-sectional area is not considered. In Fig. 5a, two linear fittings of data points from ref.(6) are demonstrated. The black dashed and blue dashed line indicate linear fittings with and without minimal cross-sectional area considered respectively. Black dashed line gives higher coefficients of determination $(R^2 = 0.98)$ than blue dashed line $(R^2 = 0.96)$. It is also clear from Fig. 5a that black dashed line can describe the beginning part of the data points better.

**Including minimal diameter in scaling law can better fit pig coronary arteriolar bifurcations**

Digitalized distribution profile from Fig. 6(B) in ref. (7) is shown in Fig. S6a. Log-normal distribution is used to fit the digitalized data as shown by the red dashed curve in Fig.S6a. We then generate 2000 random number with log normal distributions using fitted $\mu$ and $\sigma$ from Fig.S6a. Here the random number are ratio values: $(d_{\mathrm{d1}}^3 + d_{\mathrm{d2}}^3)/d_{\mathrm{m}}^3$. The minimal branch in arteriole is estimated from y-intercept of branch cross-sectional area vs. leaf number in Fig. 5b. Therefore, we have $d_0 \approx 6.2\mu m$ here. In ref.(7), only arterial vessels <50 μm in diameter are considered. The range of $d_{\mathrm{m}}$ is 6.2-50.0μm. Because the hierarchy nature of the branched network, there are more small leaf number branches than large leaf number branches. The histogram of $d_{\mathrm{m}}$ values tends to bias towards minimal diameter $d_0$. Here we use log-normal distribution to approximate the skewed distribution of mother branch diameter $d_{\mathrm{m}}$. We further assume smaller $d_{\mathrm{m}}$ values correspond to larger $(d_{\mathrm{d1}}^3 + d_{\mathrm{d2}}^3)/d_{\mathrm{m}}^3$ ratios based on our experimental observations. Based on generated values of $(d_{\mathrm{d1}}^3 + d_{\mathrm{d2}}^3)/d_{\mathrm{m}}^3$ as well as $d_{\mathrm{m}}$, we get to know $d_{\mathrm{d1}}^3 + d_{\mathrm{d2}}^3$.

In our work, we proposed a new scaling relation $d_{\mathrm{m}}^p + d_0^p = d_{\mathrm{d1}}^p + d_{\mathrm{d1}}^p$, where $p$ is the scaling exponent which depends on the network properties. From ref. (7) measurements, we assume $p = 3$. We then asked whether new scaling law can better describe the relation between two daughter branches and mother branches. The question can be rephrased as: among the two ratios: $(d_{\mathrm{d1}}^3 + d_{\mathrm{d2}}^3)/d_{\mathrm{m}}^3$ and $(d_{\mathrm{d1}}^3 + d_{\mathrm{d2}}^3)/(d_{\mathrm{m}}^3 + d_0^3)$ which one is more symmetric? Skewness is a measure of the asymmetry of the probability distribution. The definition of skewness is:

$$g_s = \frac{\sum_{i=1}^{N}(Y_i - \bar{Y})^3/N}{s^3}$$

where $\bar{Y}$ is the mean, $s$ is the standard deviation, and $N$ is the number of data points. The measured skewness of Fig. S7a is 0.91 and that of Fig. S7b is 0.29. $(d_{\mathrm{d1}}^3 + d_{\mathrm{d2}}^3)/(d_{\mathrm{m}}^3 + d_0^3)$ show smaller skewness compared with $(d_{\mathrm{d1}}^3 + d_{\mathrm{d2}}^3)/d_{\mathrm{m}}^3$.

New scaling law $d_{\mathrm{m}}^3 + d_0^3 = d_{\mathrm{d1}}^3 + d_{\mathrm{d1}}^3$ can better describe Fung's data in pig's coronary arteriolar bifurcations than traditional scaling law $d_{\mathrm{m}}^3 = d_{\mathrm{d1}}^3 + d_{\mathrm{d1}}^3$.

**Figures**

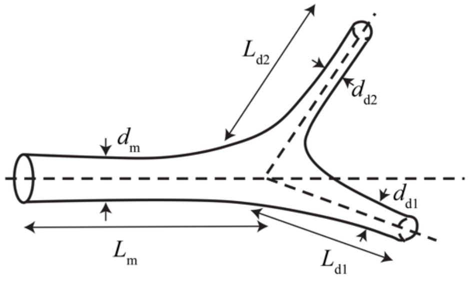


**Fig. S1 A bifurcation node.** The mother branch, with diameter $d_{\mathrm{m}}$, length $L_{\mathrm{m}}$ splits into two daughter branches with diameters $d_{\mathrm{d1}}$ and $d_{\mathrm{d2}}$, and lengths $L_{\mathrm{d1}}$ and $L_{\mathrm{d2}}$.

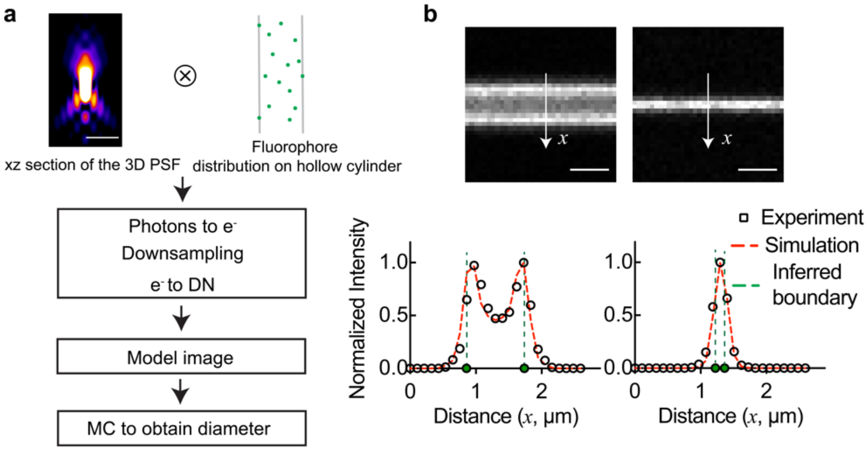


**Fig. S2 Examples of model images. a** The $x$-$z$ section of a 3D PSF is generated using the Gibson-Lanni model (8). The dendritic branch is considered a hollow cylinder randomly decorated with fluorophores. Each fluorophore is considered to be a point source and convolved with the 3D PSF. The convolved image determines the number of photons at each pixel. These photons are then transformed into a number of electrons based on quantum efficiency (QE), shot noise and the sCMOS parameters. The image is reduced to the desired camera resolution, 108 nm/pixel. These values are fed to an electron-to-DN converter (digital number, taking into account the readout noise) to obtain model images. **b** Model image examples of thick (diameter $d = 864nm$) and thin ($d = 144nm$) dendritic branches. Intensity profiles of the middle section of the two model image stacks are plotted together (black circles). Red dashed curves obtained from this super-resolution method closely track the model image measurements. The inferred diameters are 880±19 nm and 140±25 nm (mean ± SD, N=18) as shown in table I. Mean values are indicated by green dashed lines. Scale bars in (**a**) and (**b**) are 1μm.

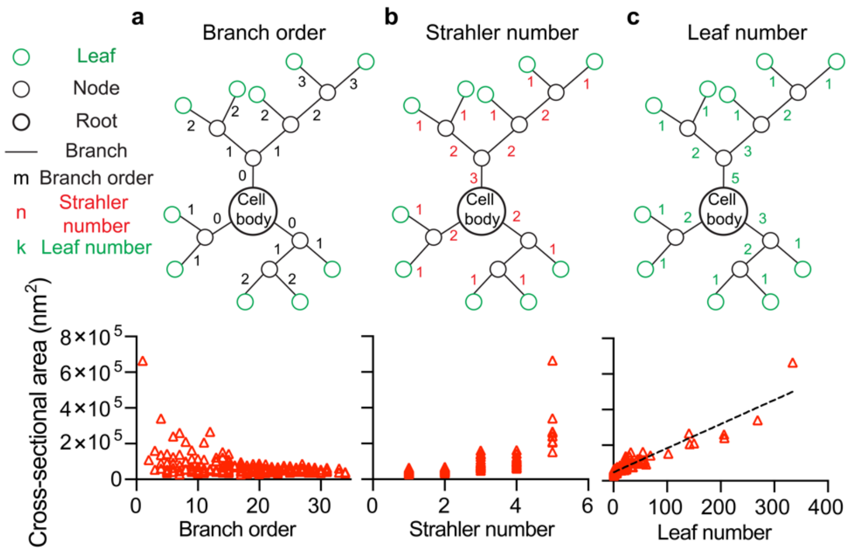


**Fig. S3 Cross-sectional area variation under different branch-ordering schemes.** Measured cross-sectional area vs. **a** Branch order; **b** Strahler number; **c** Leaf number for one 132 hr larva.

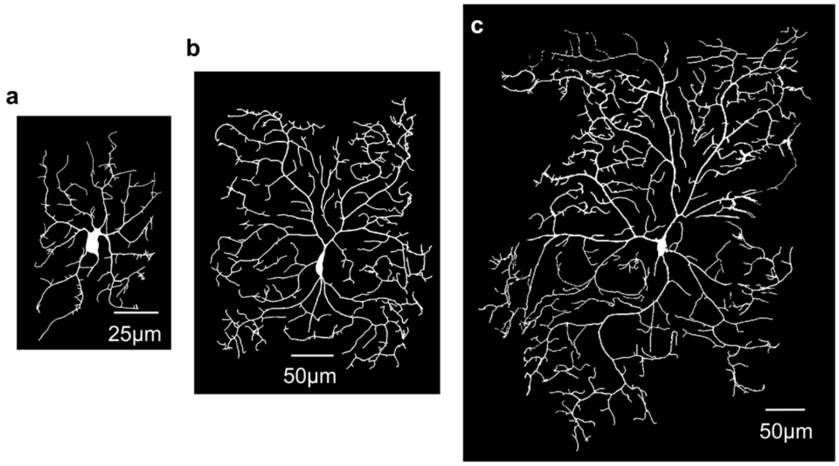


**Fig. S4 Development of Class IV neurons.** Morphologies of Class IV da neurons of age AEL of **a** 24 hr, **b** 72 hr and **c** 132 hr. Neurons were imaged using a 60X WI objective. A stitching plugin (9) was used to obtain the whole neuronal morphologies shown in **(b)** and **(c).** Class IV neurons grow throughout larval development from a width of ~60 μm at 24 hrs to ~400 μm at 132 hrs. Anterior is to the left.

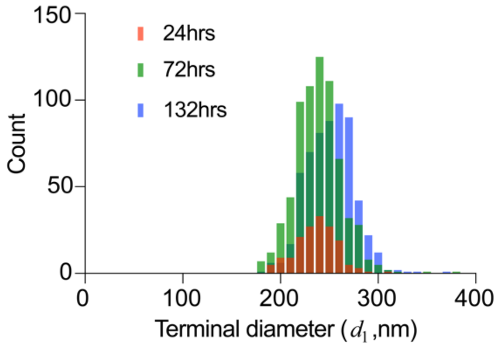


**Fig. S5 Frequency distribution of terminal diameters measured for three stages.** Six larvae were measured for each stage. Number of measurements $N$ for each stage: 24 hr ($N$=160), 72 hr ($N$ =679) and 132 hr ($N$ =598).

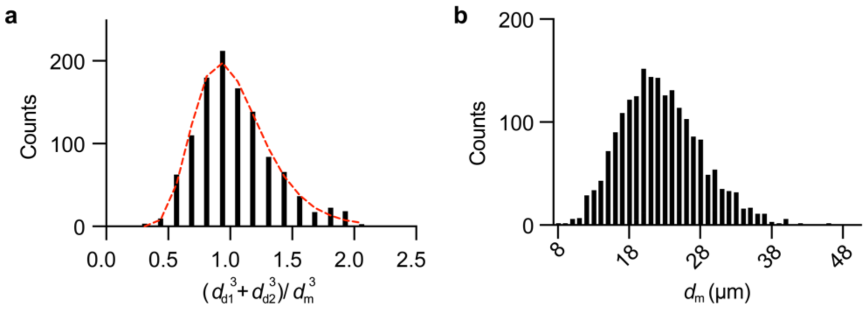


**Fig. S6 Histograms of Fig. 6B from ref. (7) and simulated mother branch distribution. a** Histogram of the ratios of sum of the cubes of daughter branch diameters to that of the mother branch for the arteriolar bifurcations. Black bars stand for digitalized data from Fig. 6B in ref. (7). Red dashed curve shows log normal fitting. **b** Log-normal distribution with $\mu = 0$ and σ=0.2 is generated and further stretched within range [8, 50] μm to simulate possible mother branch diameter $d_\mathrm{m}$ distribution.

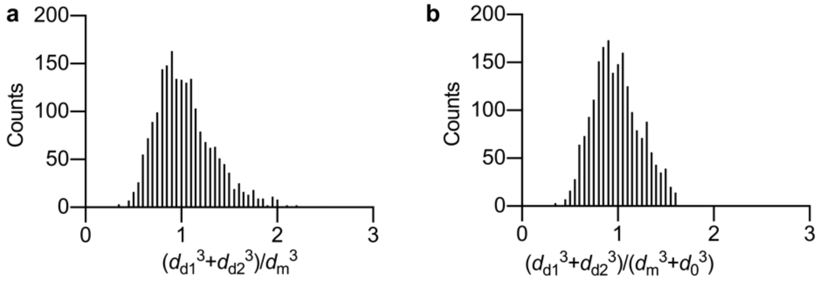


**Fig. S7 Including minimal diameter can better fit experimental data from ref.(7). a** The distribution using Eq. (2) with $p = 3$ and **b** a minimum diameter term $d_0$ of 6.2μm (estimated from y-intercept in Fig. 5b) added. The more symmetric distribution indicates a better agreement when a minimum diameter is included. Data from (7).

**SI References**


1. W. Rall, Branching dendritic trees and motoneuron membrane resistivity. *Experimental neurology* **1**, 491–527 (1959).

2. Q. Wen, D. B. Chklovskii, A Cost–Benefit Analysis of Neuronal Morphology. *J Neurophysiol* **99**, 2320–2328 (2008).

3. J. B. Hursh, CONDUCTION VELOCITY AND DIAMETER OF NERVE FIBERS. *Am J Physiology-legacy Content* **127**, 131–139 (1939).

4. D. B. Chklovskii, A. Stepanyants, Power-law for axon diameters at branch point. *Bmc Neurosci* **4**, 18 (2003).

5. A. N. Strahler, Quantitative analysis of watershed geomorphology. *Eos, Transactions American Geophysical Union* **38**, 913–920 (1957).

6. K. Shinozaki, K. Yoda, K. Hozumi, T. Kira, A quantitative analysis of plant form- the pipe model theory: I. basic analyses. *Jpn J Ecol* **14**, 97–105 (1964).

7. G. S. Kassab, Y.-C. B. Fung, The pattern of coronary arteriolar bifurcations and the uniform shear hypothesis. *Ann Biomed Eng* **23**, 13–20 (1995).

8. S. F. Gibson, F. Lanni, Experimental test of an analytical model of aberration in an oil-immersion objective lens used in three-dimensional light microscopy. *JOSA A* **9**, 154–166 (1992).

9. S. Preibisch, S. Saalfeld, P. Tomancak, Globally optimal stitching of tiled 3D microscopic image acquisitions. *Bioinformatics* **25**, 1463–1465 (2009).
